# Cancer Cell Removing Using a Reinforcement Learning Agent

**DOI:** 10.1101/2024.09.01.610680

**Authors:** Ali Mousavi Fard

## Abstract

Cancer cell is a deadly problem which is the main cause of global death. Unfortunately, the conventional therapies like chemo/radio therapy are not viable ways to remove all of the cancer cells. Although Robotic achievements have been increased in cancer therapy, these devices do not have the decision-making ability to grasp their environment like biologists. In this paper, a cancer cell removing method based on Artificial Intelligence techniques is introduced. The proposed idea adopts a combination of object detection and reinforcement model in order to detect the cancer cells and take some actions to remove them. To implement this idea, YOLOv9 is trained on a cancer cell image dataset to detect and segment the cancer cell and create a set point for RL model then in the next step, Soft Actor Critic (SAC) is considered as a RL model to grasp the desired environment and take some appropriate actions to reach the target. The experimental result of this model shows that the proposed model can be adopted in different cancer therapy robots like micro/wireless soft robots to boost their performance in terms of their decision-making ability.

## Introduction

Cancer is a deadly disease causing the majority of worldwide death. Cancer treatment has been an ambiguous issue in medical science. Nonetheless there are a wide range of research works that reveal various cancer treatments. Cancer therapy is divided into conventional techniques like surgery, chemotherapy and radiotherapy and novel approaches comprise gene therapy, immunotherapy and MSC-based therapy [1-3]. In conventional ways, the cancer tumor is extracted from patient body by adopting surgical tricks but during this operation a considerable number of cancer cells escape into blood vessels, therefore, the patient may experience metastasis. To tackle this problem a combination of treatments has been adopted to remove the survived cancer cells. After surgery, the patient undergoes treatment by using chemotherapy or radiotherapy. Chemotherapy uses chemical material such as the platinum-based chemo therapeutic to remove cancer cells, this material is injected to patient body under the doctor supervision in order to decline the cancer recurrent problems, however, it can result in adverse events and chronic morbidity because the chemical materials attack both normal and cancer cells, as a result, it can badly impact on human body and create other diseases like anemia, moreover, it weakens immune system function [1-3]. Radiotherapy adopts high energy x-rays and it aims high energy radiation beams from outside the body at the tumor to burn the cancer cells. Despite important advances in radiotherapy, there are numerous adverse effects associated with this form of treatment including diarrhea, abdominal cramps and pelvic pain, skin toxicity, lymphedema and sexual dysfunction [1-3]. Obviously, chemotherapy and radiotherapy possess high side effects and can cause diverse difficulties, consequently, researchers are striving to find an alternative for cancer treatment which do not have these drawbacks. Recently, potential of biology tricks like immunotherapy, gene therapy, and MSC-based therapy have gained abundant attentions due to their low catastrophic and side effect problems on patients. The advantage of immunotherapy is that it specifically targets and identifies the presence of abnormal cells within a tissue or organ called dysplastic precancerous. Plus, the clinical trials demonstrated that by activating and empowering the T-cells, it can provide a tumor suppressive way to remove cancer cell with body immune system [1,4-5]. Gene therapy pursues different goals to prevent the cancer progression because cancer cells proliferate with high speed. The gene therapy techniques can induce cell-cycle arrest into cancer cells, cell cycle arrest stop cancer cell division, therefore, the possibility of metastasis will be vanished. Moreover, by regulating the DNA of cancer cell using novel techniques like Crisper/Cas9 and None-coding RNA, the baleful effect of cancer tumors can be modified without invasive surgery [1]. Another insightful biology idea for cancer cell removing is MSC-based therapy. Mesenchymal stem cells (MSC) are among the most frequently used cell type for regenerative medicine. The main goal of MSCs is tissue repair and renewing the cells of whole body. MSCs also possess the capacity to migrate to injured sites in response to environmental signals and promote tissue regeneration mediated by the release of paracrine factors with pleiotropic effects [6]. MSCs are also able to inhibit the immune system, promote cell survival and induce angiogenesis among others pleiotropic activities. All these advantages make MSCs a powerful tool for clinical application in regenerative medicine. The therapeutic benefits of MSCs have prompt their use in cell-based strategies to treat different diseases, including cancer. Similar to damaged tissues, tumors exert chemoattractant provoke MSCs that influence their recruitment to tumor sites. MSCs interact with cancer cells via direct and indirect mechanisms that affect tumor development. These paracrine factors affect cellular processes involving tumor cell cycle, cell survival, angiogenesis, and immunosuppression/immunomodulation, allowing MSCs to regulate cancer by so doing, MSCs can induce cell cycle arrest and reduce cancer growth and remove them [6]. Moreover, by using MSCs and stem cells we can induce senescent into cancer cells and remove them with senescent removing drugs [7]. Despite of application of MSCs in cancer treatment, it can excite tumor development by its survival ability, therefore, the potential of MSCs in cancer therapy is controversial. Although all of the novel biology tricks are so intriguing ways for cancer cell removing, they have not reached the expected therapy level [8]. It is noteworthy that all cancer removing techniques which include conventional and novel approaches are not definitely able to remedy cancer because cancer cells gradually obtain anti-cancer drug resistance capability [8], therefore, a new intelligence technique should be geared to solve this issue with high reliability. Nowadays, application of robotic in medicine has been soared remarkably and it is widely used in diagnosis, localization and treatment with minimally invasive surgery [9]. A good example of medical robotic is endoscopy. Traditional endoscopy required open surgery and this type of operation was associated with slow patients’ recovery time, postoperative pain, increased rate and severity of postoperative complications, blood loss, and immunological stress response of the tissue. However, recent endoscopy devices by adopting robotic techniques can do this task with less invasion. These modern endoscopy devices have miniature cameras in order to provide visual feedbacks for the clinician of the internal organs of the patient. At the same time, surgical instruments can be used to probe tissue surface. Nevertheless, doctors can recognize tumors or other issues in human body and remove them in the best way [9-10]. Another advantage of medical robot is drug delivery into vital organ of patient body in special condition that the typical drug delivery ways like capsule cannot have a desired impact on damaged organ, as a result, a specific robot act as a drug delivery to transfer sufficient drug into defective site in order to repair or treat that mechanism [11]. Surgical robot which is so called medical robot that plays an essential role in diverse disease treatments such as cancer therapy and repair damaged tissue in various parts of the body. The main limitation of medical robots is their kinematic and dynamic operations and control of their positions into human bodies. To tackle this problem, soft robotic was proposed to adapt its structure with the environment of the patient body. One of the virtues of using this type of devices is that sometimes the issues are appeared in dangerous part of the body. if the doctors persist to access the expected site, patient’s life may be in danger, therefore, a device with adaptive structure has to be adopted in order to reach that point and remove the issues with minimal invasive [12]. In this field, soft robotic is named holder which is used for many medical aims. Recently, the possibility of using Micro and Nano Robot for medical tasks has been taken into account due to its speed, flexibility and tiny structure. Micro and Nano robots can easily move throughout human body and have a good performance in drug delivery or cancer cell removing. However, these devices are the new branch of robotic and it is anticipated that they will come to practical world in forthcoming future [13]. Although medical robots are so applicable and necessary for a wide variety of treatments, they have some weaknesses. The capability of artificial intelligence has not been used in these robots effectively in order to grasp their environments. Moreover, medical robots cannot able to assess and manipulate micro-scale tissue like cells, as a result, there is a research gap to create an intelligent device for cancer cell removing because for achieving this goal firstly we need a camera which can take images in micro-scale level in order to record the cell function and this camera has to be set in soft or micro robot by so doing the medical robots can manipulate cells and remove dangerous or cancerous cells in human body. Another research gap of these devices is that there is not a decision-making ability to do cancer cell removing according to its environment situation. In cancer cell removing task, the conventional treatments cannot remove all issues with high accuracy and reliability, as well as, in some situations such as Multimorbidity, the cancer cell removing must be stopped provisionally in order to treat other illnesses while conventional techniques like different drugs or operations do not have this intelligence to do it with high speed[14],therefore, micro or soft robotic must have a highly intelligence to act as a process controller in body and continue its mission until all of the cancer cells are removed. For achieving this aim, artificial intelligence techniques should be adopted in order to first detect the cancer cell then by using decision making ability take the special actions to remove these cells. In this paper, we indicate an artificial intelligence model to add the grasp of cancer cell removing skill in medical robots like biologists. This algorithm can be adopted on different medical robots to do this task. The paper idea will be discussed in next sections: in section 2, we deal with related works about application of Artificial Intelligence and Robotic in cancer therapy, in section 3, the proposed idea will be elaborated and eventually the conclusion is indicated.

### Section 2

Robotic technology has created promising performance in medical tasks such as monitoring, predict, surgical operation and treatments [15]. A good example of medical robots is endoscopy devices, these devices can enter human body by minimal invasion and take images form the expected part of patient body so that the doctors can detect the tumors or other issues [16]. In [17], the paper declared the application of endoscopy laser micro surgery, this device is transferred to human body with minimal invasion then by using its tiny camera can localize the damaged part of human body like tumor cells, as a result, doctors can remove them by using laser. This device reflects the high-power laser to burn the tumor cell or tissue [17]. The advantages of laparoscopic devices have been increased; this kind of devices can access unreached area by using its adaptive structure. Laparoscopic surgery (LS) is one of modern minimally invasive surgery (MIS) techniques which adopts small incisions and long pencil-like instruments to perform operations with a camera. Patients who choose laparoscopic surgery usually have shorter hospital stays, quicker recovery, and less post-operative complications. For improving the ability and reliability of laparoscopy, the computer vision and virtual reality techniques have been adopted in order to visualize body tissues in 3D environment by so doing surgeons can grasp the environment more effectively and are able to create a margin for cancer organs with high accuracy, as a result, in this method the cancer tumor can be more distinguishable than previous cancer therapy tasks [18]. Another use of virtual reality in laparoscopy is surgeon training. Most of the time the doctors cannot friendly interact with medical robots nevertheless training platforms should be created for doctors in order to increase their skills in using and manipulating surgical robots for real operations. Virtual reality and computer vision are the heart of these training platforms [18-19]. The MRI images can guide robotic devices for ultrasound prostate cancer therapy. The developed robotic system is guided by MRI to achieve near real-time temperature and monitor the ultrasonic exposure. It includes three linear stages and two angular stages thus offering movement of the transrectal focused ultrasound (FUS) transducer in 5 DOF. The MRI plays an operation controller in ultrasound therapy and it can inform the robotic device and specialist about progression rate of surgery or therapy [20]. Recently using novel robotic tasks such as wireless soft robotic and micro robot for cancer therapy have obtained tremendous attentions [21]. The wireless device can be transferred into body and be controlled by high power waves in order to navigate the device to accomplish its missions. Wireless soft robotic device is divided into four categories namely oral administration, Injection and insertion, Implantation, Endoscopy. Oral administration involves swallowing the device in the form of a capsule or other shapes, such as a beam shape. The device is designed to withstand the stomach acidic environment and safely traverse the gastrointestinal tract until it reaches the desired location. It can perform many functions, such as cargo delivery, surgery and sensing [21]. Injection involves directly delivering the wireless soft medical device into the blood vessels or non-vascular organs through a hypodermic needle or catheter. This method allows precise targeting of specific anatomical sites, such as connective tissues or blood vessels, and is usually used for applications such as targeted drug delivery, neurostimulation or physiological monitoring [21]. Implantation involves surgically placing the wireless soft medical device inside the body. The device may be implanted in tissues, organs or cavities to monitor physiological parameters, provide continuous therapy, or facilitate tissue regeneration. Implantation can be used to position pacemakers, neural implants or mobile scaffold structures for tissue regeneration or to assist the function of the organ [21]. Endoscopy involves the insertion of a wireless soft medical device into the body through natural orifices or small incisions using an endoscope. This method enables direct visualization and access to internal organs and cavities. Endoscopic deployment is frequently employed in gastrointestinal examinations, biopsies or minimally invasive surgeries [21]. Despite all of the virtues of wireless devices in medical tasks, they have a considerable number of limitations in fabrication and control system. The materials which are used in these devices can increase the risk of infection and inflammations in human body, therefore, the specific material should be adopted that is compatible with human organs. For example, several cancers arise in harsh environments inside the body, such as gastric cancers surrounded by acidic digestive fluid (pH 1.5–3.5) inside the stomach. Most drugs in this environment require proton pump inhibitors for neutralization. By contrast, magnesium and zinc-based chemical microrobots are perfectly adapted to this environment [22]. Microrobots have demonstrated a great potential to perform various important functions, for example, the delivery of drugs, the manipulation of the cells and biosensing through manual mechanisms [23]. These intelligent systems are proficient in information processing, signaling, sensing, actuation, and communication, as well as conducting biological tasks at the cellular level and delivering drugs locally result in improved efficacy and fewer side effects in comparison traditional therapeutics [23]. Microrobot based on bacteria, red blood cells, or stem cells are thought to be effective and compatible for the targeted delivery of drugs in body environment for various treatments like cancer cell removing. Tumors can occur at almost any site of the body, including hard to reach locations situated deep inside the body, and so-called sanctuary sites, for instance, behind the blood–brain barrier that render them invisible to conventional therapy. However, wireless soft and micro robot can detect and remove hard to reach cancer tissues with high reliability [22]. Cancer removing microrobots can be split into three major classes that can be distinguished based on their make up and source of propulsion namely cellular microrobots, synthetic microrobots and hybrid microrobots. Cellular microrobots (biologically actuated) that exclusively consist of cell-made components, and are precision engineered exhibit anticancer effects. Synthetic microrobots (chemically and/or physically actuated) contain only man-made materials, structures and components. Hybrid microrobots, consisting of both artificial and cell-made components can be propelled by biological or artificial means [22]. After reaching the tumor, microrobots need to be able to eradicate the cancer cells. Two major pathways can be distinguished comprise direct and indirect interaction. In direct interaction the microrobots attack the cancer cells and strive to eliminate them while indirect interaction tries to stimulate and empower immune system to destroy cancer cells. All of these smart devices can be adopted to be alternative to previous therapy techniques because untargeted chemo/radiotherapy can have severe side effects and it targets both normal and cancer cells. However, other obstacles in this field like highly reliable navigation and control system have remained as unsolved problems. Control and navigation systems must have the ability of dealing with the complex biological environment in human body in order to control themselves in body environment and not to deviate from their path till they reach their targets. To tackle this problem, plenty of research works are attempting to design wireless and adaptive control methods in order to enhance the controllability and commandability of these soft and miniature medical devices [21-22]. In addition to wireless and control algorithms, the competency of AI models should be taken into account to mitigate the error and weakness of control and navigation system. Artificial intelligence techniques have demonstrated their ability to assess medical in order to diagnose damaged tissues in the images like cancer tumor detection [24]. Moreover, using AI for analyzing biological disorder has been a fascinating research field in the world like inspecting biological senescent cells [25-27] or cancer cell detection in different parts of body [28-29]. Artificial intelligence can also be adopted in control task by using reinforcement learning models [30]. By using reinforcement learning an agent as a robot is created which is responsible for perceive its environment in order to do the specific tasks like pick and place or different robotic actions. Diverse robotic tasks like autonomous cars [31] or parallel robot [32] adopt reinforcement advantages to enhance their control and navigation performance. The combination of computer vision and reinforcement learning techniques can enhance controllability of the robot because in the first stage the target of set point is detected by using an object detection model then in the second stage the reinforcement learning attempts to find an optimized way for agent to reach its desired target [33]. The combination of computer vision and reinforcement models for controlling robots can be used for prevailing the control system weaknesses of microrobots and it can be considered alongside other control ideas like using remote electromagnetic waves. The sampled image of microrobot will be shown in figure below.

### Section 3

In this section, information about the proposed idea and the dataset used will be elaborated.

#### 3.1. Proposed Idea

According to Fig.1, in this paper, we reveal a new model for automatic control micro or soft robots in body. Our idea considers a combination of instance segmentation and reinforcement model. The aim of this idea is that in the first stage, the robot agent detects the targeted cancer cells with high accuracy and extracts the edge of the target object then by using reinforcement learning model, we can take a set of actions that help robot agent to reach its target and remove it. In this research work, we focus on cancer cell removing, nonetheless, in the first stage an instance segmentation model should be trained on a dataset which contains 900 cancer cell images. To do this task, YOLOV9 [34] is adopted to detect and segment the tumor on images with high accuracy and a public cancer cell image dataset is also considered [35]. The object detection model was trained on this cancer image dataset with 100 epochs due to GPU limitations. The result of object detection model is shown in figure below.

**Fig. 1.**
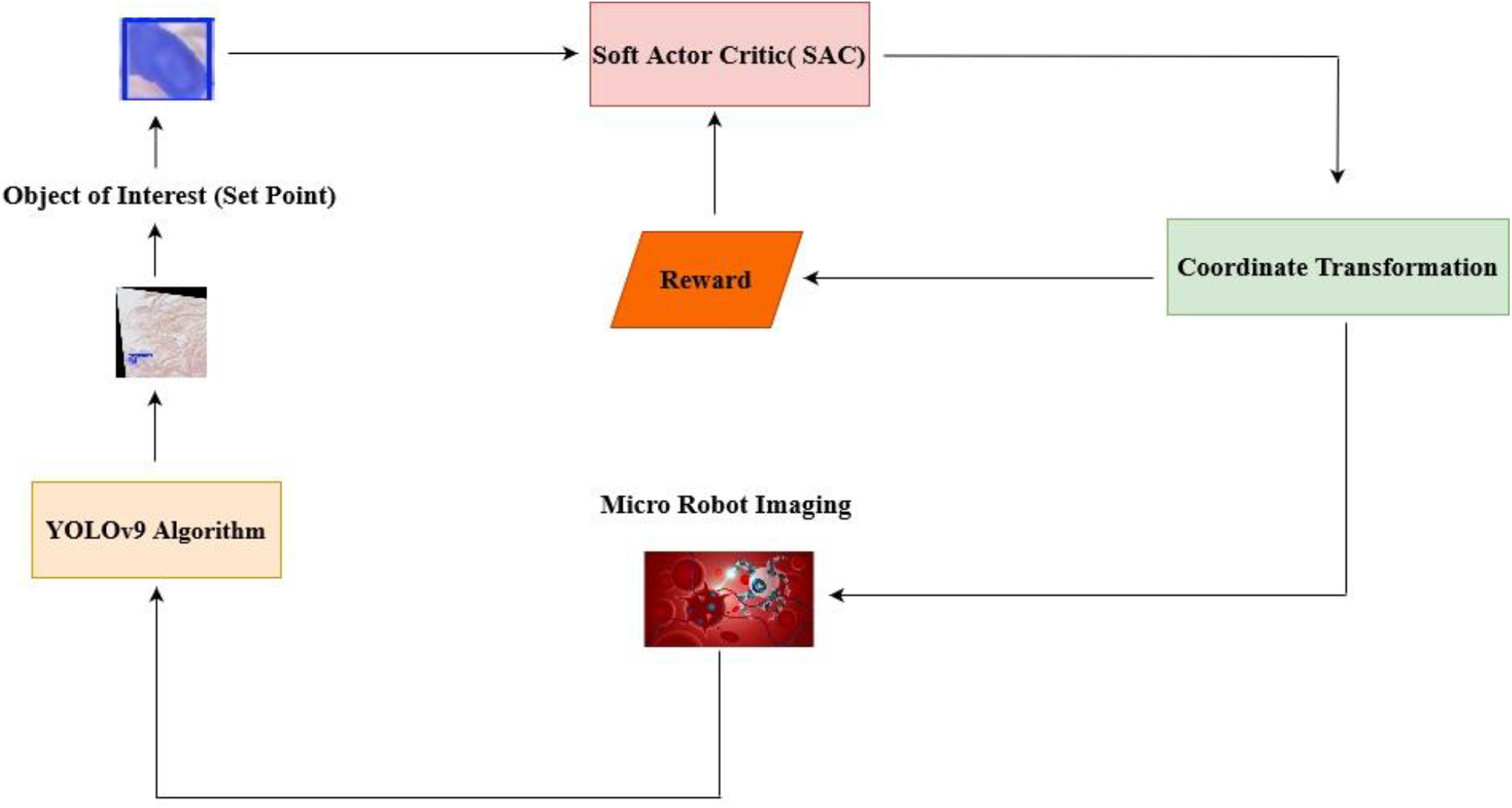
The proposed Idea.

**Fig. 2.**
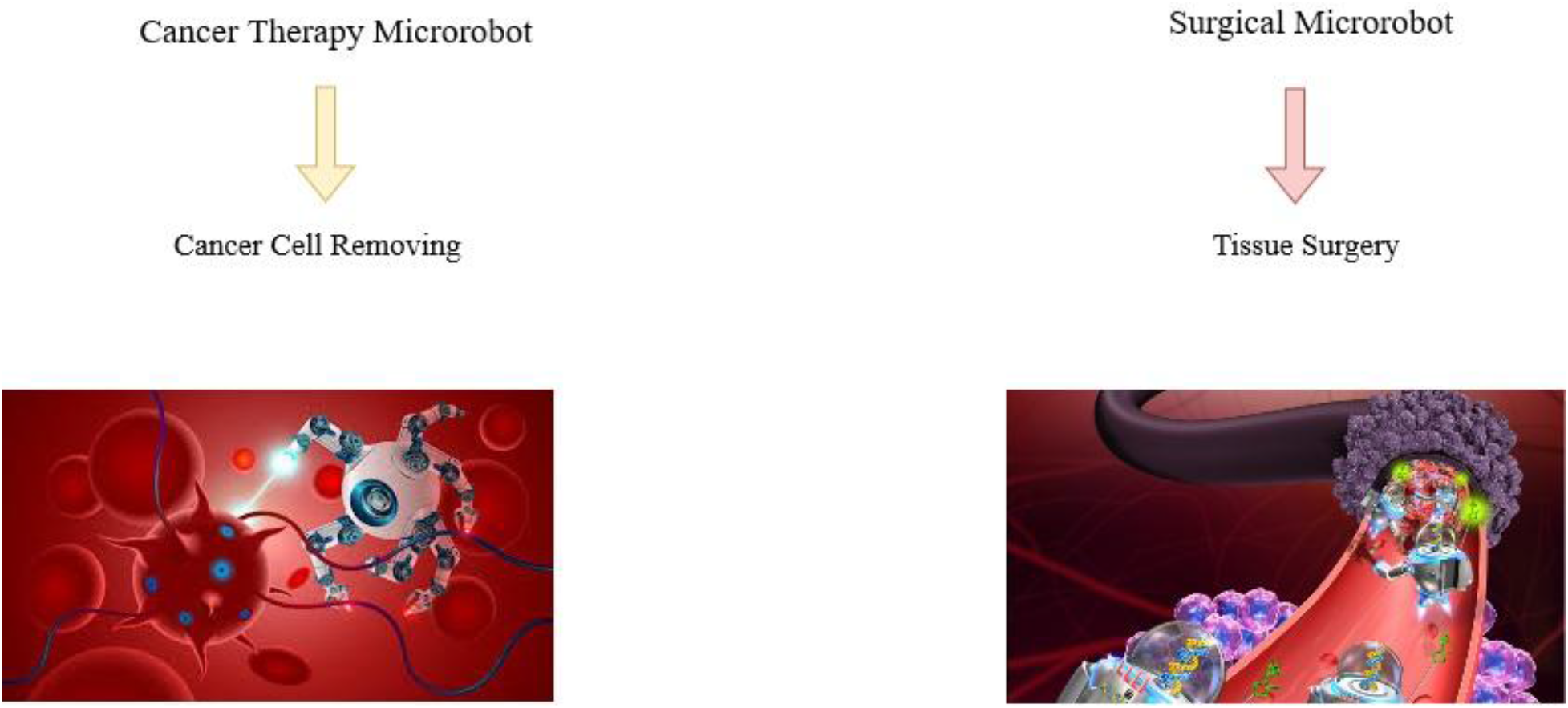
Application of Microrobot.

According to Fig.3 the YOLOv9 can detect and segment the cancer cells with high accuracy. In order to apply reinforcement learning model for cancer cell removing model, a set point or target should be assigned for RL agent, as a result, the region of interest must be extracted in image. In this researchو the region of interest or target is cancer cell and agent must learn how to reach it. To do this task, the set point must be extracted from output image of YOLOv9 model then the set point region should be cropped to create an appropriate set point for RL agent, then in the second stage the cropped image is transferred to the input of reinforcement model to inform the agent about target. The set point creation will be delineated in Fig.4.

**Fig. 3.**
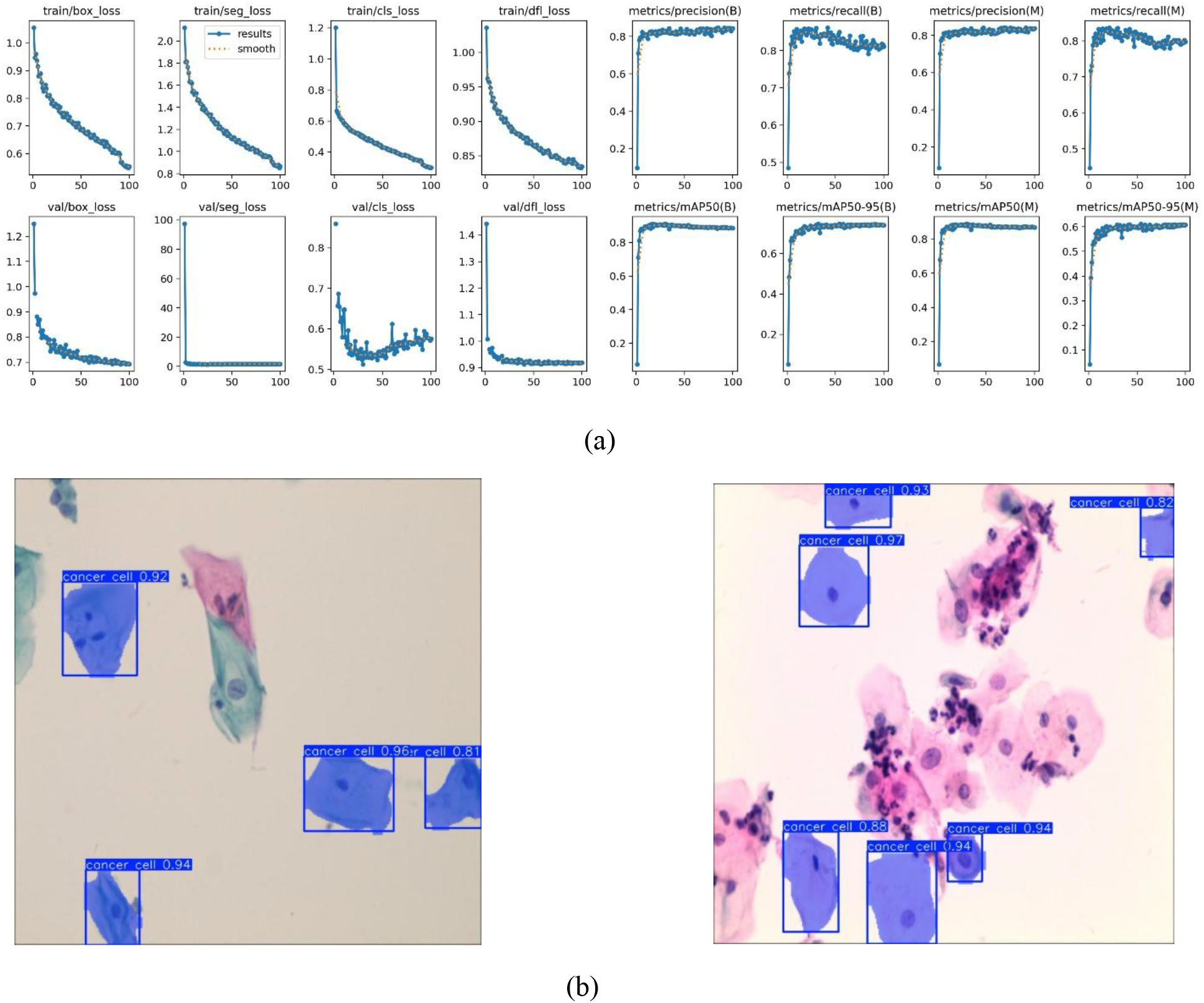
The result of YOLOv9 instance segmentation, Fig.3.a is the training result, Fig.3.b is the output result of YOLOv9.

**Fig. 4.**
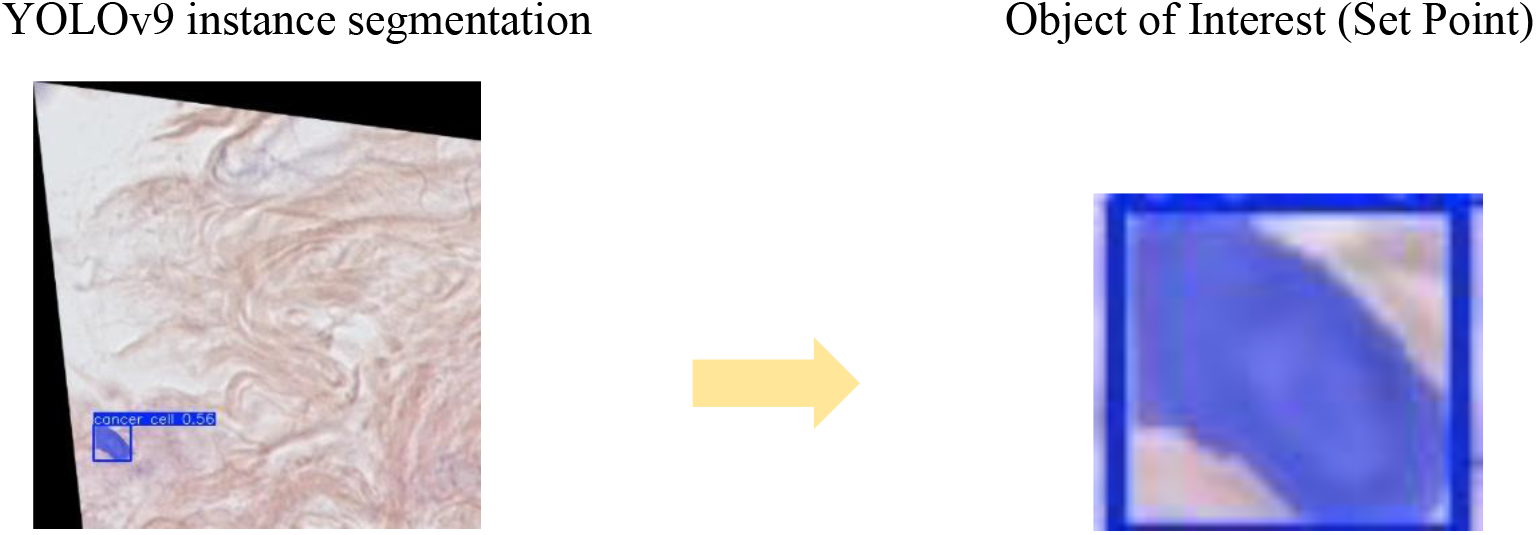
Set Point Creation for Reinforcement Model.

In second stage, a suitable Reinforcement learning must be adopted to learn the desired environment effectively. For this task, we designed simple 2D environment by using Pygame and OpenCV in order to complete the primary test of the proposed model. In this research work, we used soft actor critic model (SAC) [36]. SAC is a deep reinforcement learning algorithm that can enable a robot to learn in the real world. SAC has four features including :1) it is based on the Actor-Critic framework; 2) it can learn based on past experience, off-policy, to achieve improved efficiency in sample usage; 3) it belongs to the category of Maximum Entropy Reinforcement Learning and can improve stability and exploration; and 4) it requires fewer parameters [37]. In particular, the policy function and the soft action-value function are the actor and the critic in the Actor-Critic framework, respectively. Under state *s*, the soft action-value function will output the expected reward for selecting action *a*, thus guiding the policy function to learn. Based on the current state, the policy function will output an action to yield the system state for the next moment. By repeating these procedures, one can collect past experience to be used in training the soft action-value function. Since SAC is a random policy, the outputs of SAC are therefore the mean and standard deviation of probability distribution of the action space. Then an optimizer is adopted to update and optimize the action value in order to reach optimized way for completing the task. In this research, SAC is trained on mentioned simple environment to learn the desired space then the learned model is used in proposed idea model. The experimental result of SAC model is shown in Fig.5.

**Fig. 5.**
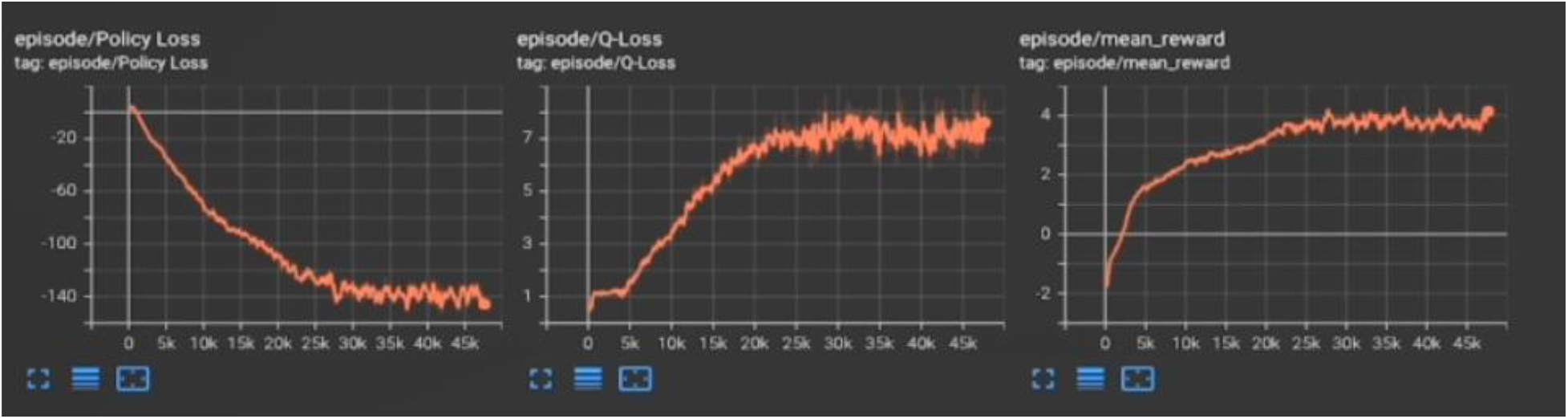
Soft Actor Critic (SAC) Result.

After training Reinforcement model, the trained RL model can be used in cancer cell removing model in order to give a target image as an input image then take some actions to reach the cancer. The proposed structure can gain a viable control algorithm for micro or wireless soft robots [21]. It can be concluded that by using a set of control method like Wireless techniques and Reinforcement based model, we can provide an avenue for enhancing reliability of these intelligent devices for adopting in diverse medical cancer treatments and other surgical operations.

## Conclusion

In this paper, an artificial intelligent technique for micro/ wireless soft robot was indicated. As the previous methods are not able to boost the intelligent ability of microrobot to grasp its environment, the proposed idea attempted to add decision making capability to this type of intelligent devices in order to detect the target with high accuracy then by using their decision-making ability, they can take some actions to reach the target. This research work dealt with cancer cell removing and a combination of object detection and reinforcement learning model was introduced as a proposed method. In the first step of the proposed idea, the desired object must be detected in the image nonetheless, a cancer cell image dataset with almost 800 samples was adopted. We trained YOLOv9 instance segmentation on this dataset with 100 epochs as a result YOLOv9 could detect and segment the cancer cell with high accuracy then in the second step, the object of interest (set point) had to be extracted from detected image, as a result, the detected image was cropped and set point was gained then Soft Actor Critic (SAC) was considered as a reinforcement learning model and was trained on a simple custom environment which was similar to real micro/ wireless soft robot environment. Eventually, these models were assembled to create the suggested model. This proposed model is able to detect the cancer cell with high accuracy then by using its grasp it takes some actions to remove them like a biologist. This technique can be adopted for any robots to do different medical treatments. For future research work, we can focus on reinforcement learning model of this method and design a reinforcement model which is able to learn the desired environment in order to boost the reliability factor of autonomous micro/wireless soft robot.

## Reference

1. Burmeister CA, Khan SF, Schäfer G, Mbatani N, Adams T, Moodley J, Prince S. Cervical cancer therapies: Current challenges and future perspectives. Tumour Virus Research. 2022 Jun 1; 13:200238.

2. Debela DT, Muzazu SG, Heraro KD, Ndalama MT, Mesele BW, Haile DC, Kitui SK, Manyazewal T. New approaches and procedures for cancer treatment: Current perspectives. SAGE open medicine. 2021 Aug; 9:20503121211034366.

3. Ghasemi Darestani N, Gilmanova AI, Al-Gazally ME, Zekiy AO, Ansari MJ, Zabibah RS, Jawad MA, Al-Shalah SA, Rizaev JA, Alnassar YS, Mohammed NM. Mesenchymal stem cell-released oncolytic virus: an innovative strategy for cancer treatment. Cell Communication and Signaling. 2023 Feb 24;21(1):43.

4. Verma J, Warsame C, Seenivasagam RK, Katiyar NK, Aleem E, Goel S. Nanoparticle-mediated cancer cell therapy: Basic science to clinical applications. Cancer and Metastasis Reviews. 2023 Sep;42(3):601–27.

5. Fuentes-Antrás J, Guevara-Hoyer K, Baliu-Piqué M, García-Sáenz JA, Pérez-Segura P, Pandiella A, Ocaña A. Adoptive cell therapy in breast cancer: a current perspective of next-generation medicine. Frontiers in Oncology. 2020 Oct 27; 10:605633.

6. Hmadcha A, Martin-Montalvo A, Gauthier BR, Soria B, Capilla-Gonzalez V. Therapeutic potential of mesenchymal stem cells for cancer therapy. Frontiers in bioengineering and biotechnology. 2020 Feb 5; 8:43.

7. Wang L, Lankhorst L, Bernards R. Exploiting senescence for the treatment of cancer. Nature Reviews Cancer. 2022 Jun;22(6):340–55.

8. Debela DT, Muzazu SG, Heraro KD, Ndalama MT, Mesele BW, Haile DC, Kitui SK, Manyazewal T. New approaches and procedures for cancer treatment: Current perspectives. SAGE open medicine. 2021 Aug; 9:20503121211034366.

9. Abad SA, Arezzo A, Homer-Vanniasinkam S, Wurdemann HA. Soft robotic systems for endoscopic interventions. In Endorobotics 2022 Jan 1 (pp. 61–93). Academic Press.

10. Li Y, Peine J, Mencattelli M, Wang J, Ha J, Dupont PE. A soft robotic balloon endoscope for airway procedures. Soft Robotics. 2022 Oct 1;9(5):1014–29.

11. Joyee EB, Pan Y. Additive manufacturing of multi-material soft robot for on-demand drug delivery applications. Journal of Manufacturing Processes. 2020 Aug 1; 56:1178–84.

12. Dawood AB, Fras J, Aljaber F, Mintz Y, Arezzo A, Godaba H, Althoefer K. Fusing dexterity and perception for soft robot-assisted minimally invasive surgery: What we learnt from STIFF-FLOP. Applied Sciences. 2021 Jul 17;11(14):6586.

13. Koleoso M, Feng X, Xue Y, Li Q, Munshi T, Chen X. Micro/nanoscale magnetic robots for biomedical applications. Materials Today Bio. 2020 Sep 1; 8:100085.

14. Cesario A, D’Oria M, Calvani R, Picca A, Pietragalla A, Lorusso D, Daniele G, Lohmeyer FM, Boldrini L, Valentini V, Bernabei R. The role of artificial intelligence in managing multimorbidity and cancer. Journal of Personalized Medicine. 2021 Apr 19;11(4):314.

15. Bramhe S, Pathak SS. Robotic surgery: a narrative review. Cureus. 2022 Sep;14(9).

16. Chauhan M, Chandler JH, Jha A, Subramaniam V, Obstein KL, Valdastri P. An origami-based soft robotic actuator for upper gastrointestinal endoscopic applications. Frontiers in Robotics and AI. 2021 May 10; 8:664720.

17. Mattos LS, Acemoglu A, Geraldes A, Laborai A, Schoob A, Tamadazte B, Davies B, Wacogne B, Pieralli C, Barbalata C, Caldwell DG. μRALP and beyond: Micro-technologies and systems for robot-assisted endoscopic laser microsurgery. Frontiers in Robotics and AI. 2021 Sep 8; 8:664655.

18. Sui Y, Pan JJ, Qin H, Liu H, Lu Y. Real-time simulation of soft tissue deformation and electrocautery procedures in laparoscopic rectal cancer radical surgery. The International Journal of Medical Robotics and Computer Assisted Surgery. 2017 Dec;13(4): e1827.

19. Privitera L, Paraboschi I, Cross K, Giuliani S. Above and beyond robotic surgery and 3D modelling in paediatric cancer surgery. Frontiers in Pediatrics. 2021 Dec 20; 9:777840.

20. Giannakou M, Drakos T, Menikou G, Evripidou N, Filippou A, Spanoudes K, Ioannou L, Damianou C. Magnetic resonance image–guided focused ultrasound robotic system for transrectal prostate cancer therapy. The International Journal of Medical Robotics and Computer Assisted Surgery. 2021 Jun;17(3): e2237.

21. Wang T, Wu Y, Yildiz E, Kanyas S, Sitti M. Clinical translation of wireless soft robotic medical devices. Nature Reviews Bioengineering. 2024 Mar 11:1–6.

22. Schmidt CK, Medina-Sánchez M, Edmondson RJ, Schmidt OG. Engineering microrobots for targeted cancer therapies from a medical perspective. Nature Communications. 2020 Nov 5;11(1):5618.

23. Suhail M, Khan A, Rahim MA, Naeem A, Fahad M, Badshah SF, Jabar A, Janakiraman AK. Micro and nanorobot-based drug delivery: an overview. Journal of Drug Targeting. 2022 Apr 21;30(4):349–58.

24. Majib MS, Rahman MM, Sazzad TS, Khan NI, Dey SK. Vgg-scnet: A vgg net-based deep learning framework for brain tumor detection on mri images. IEEE Access. 2021 Aug 18; 9:116942–52.

25. He L, Li M, Wang X, Wu X, Yue G, Wang T, Zhou Y, Lei B, Zhou G. Morphology-based deep learning enables accurate detection of senescence in mesenchymal stem cell cultures. BMC biology. 2024 Jan 2;22(1):1.

26. Kusumoto D, Seki T, Sawada H, Kunitomi A, Katsuki T, Kimura M, Ito S, Komuro J, Hashimoto H, Fukuda K, Yuasa S. Anti-senescent drug screening by deep learning-based morphology senescence scoring. Nature communications. 2021 Jan 11;12(1):257.

27. Duran I, Pombo J, Sun B, Gallage S, Kudo H, McHugh D, Bousset L, Barragan Avila JE, Forlano R, Manousou P, Heikenwalder M. Detection of senescence using machine learning algorithms based on nuclear features. Nature Communications. 2024 Feb 3;15(1):1041.

28. He W, Liu T, Han Y, Ming W, Du J, Liu Y, Yang Y, Wang L, Jiang Z, Wang Y, Yuan J. A review: The detection of cancer cells in histopathology based on machine vision. Computers in Biology and Medicine. 2022 Jul 1; 146:105636.

29. Dar RA, Rasool M, Assad A. Breast cancer detection using deep learning: Datasets, methods, and challenges ahead. Computers in biology and medicine. 2022 Oct 1; 149:106073.

30. Brunke L, Greeff M, Hall AW, Yuan Z, Zhou S, Panerati J, Schoellig AP. Safe learning in robotics: From learning-based control to safe reinforcement learning. Annual Review of Control, Robotics, and Autonomous Systems. 2022 May 3;5(1):411–44.

31. Kiran BR, Sobh I, Talpaert V, Mannion P, Al Sallab AA, Yogamani S, Pérez P. Deep reinforcement learning for autonomous driving: A survey. IEEE Transactions on Intelligent Transportation Systems. 2021 Feb 9;23(6):4909–26.

32. Lu Y, Wu C, Yao W, Sun G, Liu J, Wu L. Deep reinforcement learning control of fully-constrained cable-driven parallel robots. IEEE Transactions on Industrial Electronics. 2022 Sep 9;70(7):7194–204.

33. Taghibakhshi A, Ogden N, West M. Local navigation and docking of an autonomous robot mower using reinforcement learning and computer vision. In2021 13th International Conference on Computer and Automation Engineering (ICCAE) 2021 Mar 20 (pp. 10–14). IEEE.

34. Wang CY, Yeh IH, Liao HY. YOLOv9: Learning what you want to learn using programmable gradient information. arXiv 2024. arXiv preprint 2402.13616.

35. https://universe.roboflow.com/trialplace/cancer-cell.

36. Haarnoja T, Zhou A, Abbeel P, Levine S. Soft actor-critic: Off-policy maximum entropy deep reinforcement learning with a stochastic actor. InInternational conference on machine learning 2018 Jul 3 (pp. 1861–1870). PMLR.

37. Chen YL, Cai YR, Cheng MY. Vision-based robotic object grasping—a deep reinforcement learning approach. Machines. 2023 Feb 12;11(2):275.

